# Health workers’ perspectives on the outcomes, enablers, and barriers to the implementation of HIV “Treat All” guidelines in Abuja Nigeria

**DOI:** 10.1101/523548

**Authors:** Solomon Odafe, Kristen A. Stafford, Aliyu Gambo, Dennis Onotu, Mahesh Swaminathan, Ibrahim Dalhatu, Uzoma Ene, Ademola Oladipo, Ahmed Mukhtar, Ramat Ibrahim, Akipu Ehoche, Henry Debem, Andrew T. Boyd, Sunday Aboje, Bola Gobir, Manhattan Charurat

## Abstract

**Introduction:** To improve access to lifesaving treatment for all people living with HIV (PLHIV), Nigeria implemented the Treat All guidelines in 2016. However, health workers’ perspectives on the implementation of the guidelines have not been evaluated.

**Methods:** We conducted in-depth interviews to explore health workers’ perspectives on the Treat All guidelines. Using purposive sampling, 20 health workers providing HIV patient care were recruited from six primary health care hospitals in Abuja to participate in semi-structured interviews. Data exploration was conducted using thematic content analysis.

**Results:** The five main themes that emerged were (1) the perceived benefits of guidelines use, (2) the perceived disadvantages of using the guidelines, (3) reported patients’ response to guideline change, (4) perceived barriers/enablers to guideline use and (5) health workers’ recommendations for improvement. Health workers perceived that the implementation of the Treat All guidelines has considerably improved patient care, particularly in increasing access to skilled health care, reducing stress on patients, and increasing hope for a better health outcome among patients. Other perceived benefits were reduced mortality, reduced pre-treatment attrition, reduction in delays between case detection and initiation on treatment. Perceived major disadvantages were increased workload and overcrowded clinics. Health workers reported that most patients were willing to start treatment early. Perceived key factors enabling guidelines use were health workers’ interest, patient benefits, training and availability of tools for implementation of guidelines, government supervisory visits and hospital management support. Perceived key barriers were poverty, inadequate human resources, lack of familiarity with guidelines, and lack of consistent supply of HIV test kits at some sites.

**Conclusions:** Implementation of the Treat All guidelines led to perceived improvement in patient care. Further improvements could be achieved by implementing an efficient supply chain system for HIV rapid test kits, and through guidelines distribution and training. Additionally, implementing differentiated approaches that decongest clinics, and programs that economically empower patients could improve access to treatment.

## Introduction

HIV/AIDS continues to be a major public health disease accounting for 35 million deaths across the world [1]. In 2016 alone, there were 1.8 million new HIV infections and 1 million deaths worldwide [1]. At the end of 2016, there were approximately 36.7 million people living with HIV (PLHIV) including 2.1 million children worldwide [1]. Since the introduction of antiretroviral therapy (ART), there has been a 43% reduction in annual deaths due to HIV/AIDS globally [2]. By mid-2017, there were approximately 20.9 million PLHIV including 120,000 children receiving lifesaving ART; this number represents only 57% of the total PLHIV globally [3]. Africa remains the worst hit continent by HIV/AIDS, accounting for 25.6 million (70%) of PLHIV globally [1]. The United Nations General Assembly Special Session 2001 (UNGASS-HIV) has committed to ending the AIDS epidemic by 2030 [2]. In June 2016, UNGASS signed on to a political declaration of the 90-90-90 strategy, with the objectives to identify 90% of individuals who are HIV positive, treat 90% of all identified individuals who are HIV positive and achieve viral suppression in 90% of those on treatment by 2020 [4]. A key strategy for achieving epidemic control is increasing access to lifelong ART [2]. In order to improve access to ART, in June 2016, the World Health Organization launched the implementation of the “Treat All” guidelines, which recommend the immediate treatment of all individuals who test positive for HIV [1].

The Federal Republic of Nigeria is located in West Africa and has a population of approximately 191 million people [5]. The country has the second largest HIV burden in the world with estimated 3.1 million PLHIV [6]. In 2017, there were about 210,000 new cases of HIV infections and about 150,000 new AIDS- associated deaths in Nigeria [6]. In 2017, an estimated 1,039,520 PLHIV were receiving ART in the country [6]; thus, only one-third of the persons living with HIV in the country received ART [6]. Subsequent to the release of the UNAIDS treatment target within the 90-90-90 strategy, the President’s Emergency Plan for AIDS Relief (PEPFAR) in Nigeria piloted the Treat All approach in clinical management of PLHIV in 32 local government areas (LGAs) between March 2016 and December 2016 [7]. In December 2016, the government of Nigeria, through its Federal Ministry of Health (FMOH), published the guidelines to treat all individuals testing HIV positive in the country to help improve access to lifesaving ART [8]. However, there are currently shortages of human resources across all cadres of health staff in Nigeria due to increasing migration of health care workers to areas offering higher compensation, poorly motivated health workforce, and mal-distribution of health staff [9]. Consequently, there have been concerns that the new HIV Treat All guidelines will increase the burden on an already overstretched health system and potentially compromise care for PLHIV. Additionally, since HIV treatment program managers plan to scale up implementation of the Treat All approach nationally, knowing health workers’ perspective on potential enablers and barriers to implementation is necessary for devising effective implementation strategies. Thus, this study aimed to understand health workers’ perspectives on the implementation of the Treat All guidelines for HIV treatment in Nigeria, specifically their perceptions of the guidelines’ impact on the quality of patient care, outcomes for PLHIV, and of potential enablers and barriers of effective guideline implementation.

## Materials and Methods

### Study Design

This was a qualitative study of health workers’ perceptions using an in-depth interview approach. We utilized an interpretivist epistemological approach, reinforced by a social constructivist standpoint, to obtain the perspective of participants on the research topic [10]. By using this approach, researchers were able to explore participants’ understanding and perceptions on the implementation of Treat All guidelines at their various clinics [10].

### Study setting

The study was conducted in six health facilities in Abuja Municipal Area Council in the Federal Capital Territory (FCT), Abuja, Nigeria. Abuja is the administrative capital of Nigeria and has a projected population of 3.56 million [11]. The FCT Abuja has approximately 80,000 PLHIV with an HIV prevalence of 5.8% [12]. As of June 2017, there were approximately 43,000 HIV-positive individuals receiving ART in FCT Abuja [13].

### Sampling and inclusion/exclusion criteria

Health worker interviews were conducted at six hospitals between February and March 2018 within LGAs in FCT Abuja where the Treat All guidelines had been piloted between March and December 2016 [7]. Health workers were purposefully selected for interview [14]. Study population inclusion criteria included 1) health worker, who may have been a doctor, pharmacist, nurse, laboratory scientist, record officer, or data analyst or program manager; 2) working at the HIV treatment program and 3) has knowledge about the HIV Care and Treatment program, determined by a minimum of three years of experience working in the HIV program. All individuals who either did not meet inclusion criteria or refused to give consent were excluded from study.

### Data collection methods

All interviews were conducted at the study site, in environments familiar to the participants. The researchers sought permission from each participant before digitally recording interview sessions. Additionally, a record of important statements made or activities observed during the fieldwork was kept as notes from the sessions.

### Study Instrument

The study used a specially developed interview guide based on a previous study which evaluated guidelines among health workers in Kenya [15]. All interviews were conducted in English. The interview guide covered knowledge about the health workers’ experience using the Treat All guidelines, the care and outcomes of patients since the introduction of the Treat All guidelines, and perceived enablers or barriers to using the guidelines in clinical care. The interview guide was piloted with three health workers each from one of three different departments at a comprehensive HIV treatment center in Abuja.

### Data Analysis

The researchers utilized thematic content analysis to explore the data collected during the interviews [16]. After the interview, participants’ responses were transcribed verbatim and the lines of text numbered. All transcripts were anonymized to ensure confidentiality. The transcription process was followed by coding of the transcripts. One qualitative coder manually reviewed all transcripts. An inductive approach in which codes emerged from the content of the raw data rather than a preconceived theory was used during data analysis [17]. In addition, respondent validation and peer review were utilized to avoid coding bias [18]. Initial descriptive codes created after review of all transcripts were recoded into analytic codes based on emerging themes from all transcripts. A codebook containing both descriptive and analytic codes that related major themes and sub-themes guided the development of the study discussion and conclusion. To ensure trustworthiness, after transcribing audio recordings, the researchers reviewed transcripts with respondents to ensure the content captured what participants intended and used appropriate quotes during reporting [18].

### Ethical consideration

Ethical approvals were obtained from the National Health Research Ethics Committee of Nigeria and the U.S. Centers for Disease Control and Prevention institutional review board (IRB). An informed consent was obtained from all participants interviewed. Information provided during the interview did not include personally identifiable information and only aggregated data and unlinked quotes were used for the study reports. All participants’ names were replaced with numbers in the transcripts, during analysis and reporting [19].

### Results

A total of 20 health workers were interviewed in the study, comprising 8 medical doctors, 5 nurses/ counsellors, 4 laboratory scientists, 2 monitoring and evaluation (M&E) officers and one data analyst. From the qualitative analysis of their interview responses, we identified five major themes of importance describing (1) the perceived benefits of guidelines use, (2) the perceived disadvantages of using the guidelines, (3) reported patients’ response to guidelines change, (4) perceived barriers/enablers to guidelines use and (5) health workers’ recommendations for improvement. Table 1 summarizes the themes, sub-themes, and codes from the analysis of the transcripts from the interviews. These themes and sub-themes were then reorganized into principles within four domains: perceived benefits of the change in guidelines on clinical practice, reported patients’ response to guidelines change, perceived patient outcomes with the guidelines, and factors affecting guidelines use.

**Table 1:**
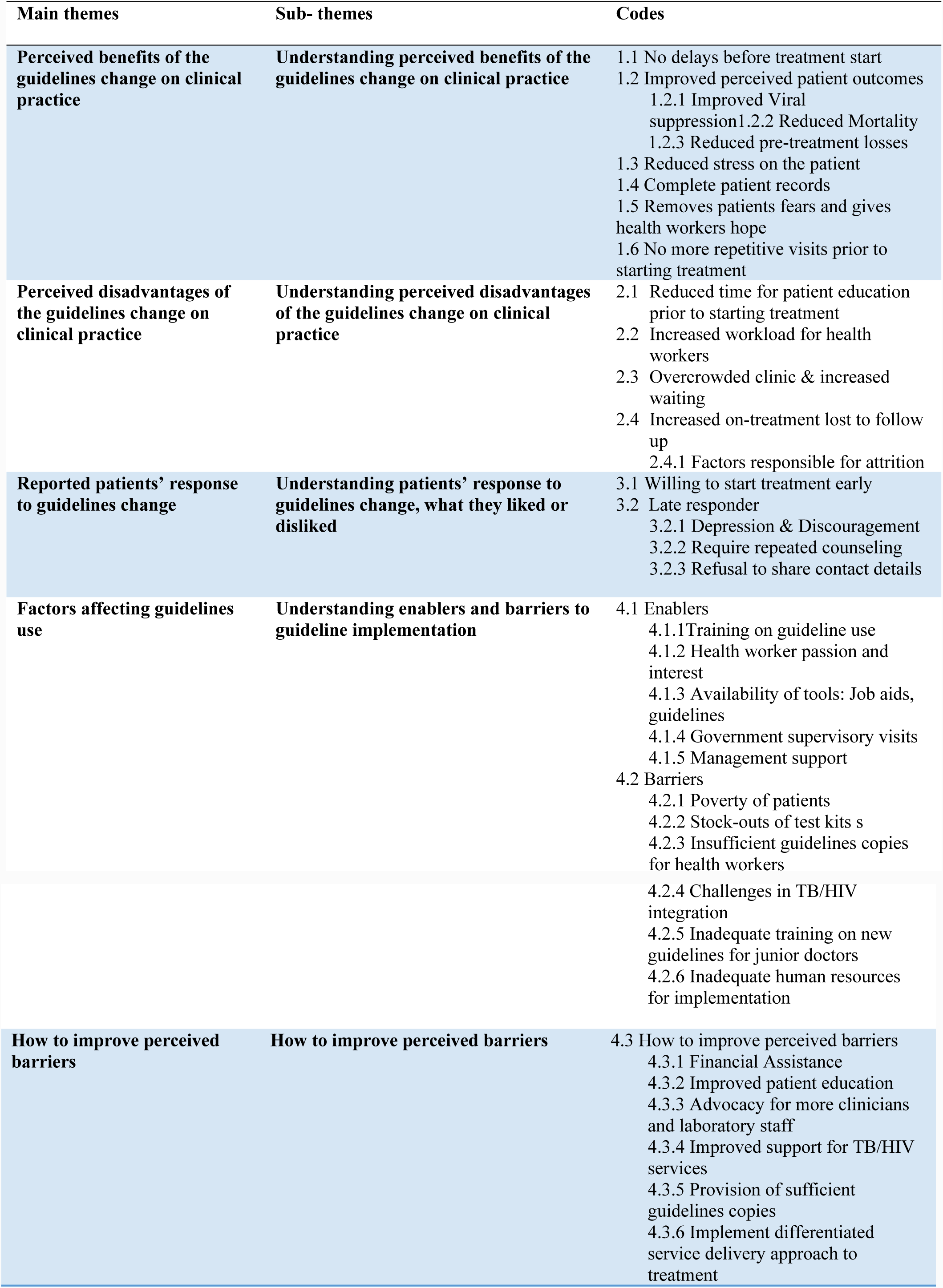
Coding scheme from thematic content analysis

### The perceived benefits of the Treat All guidelines

Perceived benefits experienced due to the introduction of the new national Treat All guidelines were described at two broad levels, benefits to the patients and benefits to the health worker or the health system. The following sections describes the benefits of the Treat All guidelines to these broad stakeholders.

#### The perceived benefits of the Treat All guidelines to the patients

The majority of health workers interviewed in the study believed that the Treat All approach has been of tremendous help in improving patient care. The most important benefit described was early initiation on ART. Staring treatment early means doctors no longer wait until patients’ health deteriorates before commencing ART. Furthermore, health workers also perceived that the new approach may have helped in retaining more persons previously tested HIV positive but waiting to be started on treatment:

*(Counsellor):“It is helping also to initiate treatment early and of course help to retain patients in care before starting treatment. In the past, we delayed patient treatment and they go away”*

Additionally, starting treatment immediately after testing means the need for repetitive follow-up visits, delay in treatment start and risk of losing the patient between the time of test and time of treatment initiation are reduced.

*(Counsellor):“we take them round and round, they go and come before they receive their treatment but with the test and treat [Treat All] we are able to give them the treatment and they go and they are happy for it”*

*(Clinician):“The initial loss after diagnosis has greatly reduced. You know the way our society is structured, sometimes when they go they never come back. They end up in the churches and some in other places.”*

Another important benefit of the Treat All approach was a perception of improvement in patients’ health outcomes. The majority of respondents reported that since the start of the guidelines, there have been increased testimonies from patients that they were doing much better:

*(Clinician):“The testimony we have is some of them returning to say (…) Doctor since we started this drug I am getting much better”*

*(Clinician): There has been great advantage, because it is a disease we cannot cure. But, if you are starting treatment immediately it makes the patient live healthier and longer”*

Moreover, the majority of the participants reported that among patients who had their viral load test done, they observed an improved viral suppression rate since the implementation of the Treat All guidelines. One perceived reason for the improvement was that patients do not wait until they are very sick before starting ART:

*(Clinician): “By the time we do their viral load test after six months we find out that they are virologically suppressed. (…) I will see those with test and treat [Treat All] are actually doing better”*

Consequently, there has been a perceived reduction in the incidence of opportunistic infections (OIs) and mortality:

*(Clinician): “(…) you tend not to have more OIs, mortality is being reduced, and transmission rates also reduce”*

Additionally, the majority of the health workers believed that there have been mostly HIV negative babies born to HIV infected mothers since the implementation of the new guidelines:

*(Clinician): “We have not had any positive child since we started implementing the guideline[s] (….)”* A common perception among respondents was that the implementation of the previous HIV treatment guidelines was very stressful for the patients. It required multiple clinic visits, frequent blood draws for CD4 investigation, and long hours of repeated counselling sessions before treatment was initiated. The process required multiple interactions between patients and health workers, some of which could be perceived as unnecessary, time wasting and stressful. However, the implementation of the Treat All guidelines has simplified the process and reduced the stress on the patient. Furthermore, participants explained that HIV patients’ frequent visits to the clinic for drug pick up usually raises questions in the community. Community members often want to know why patients visit the clinics so frequently. Some patients stayed away from clinic to avoid being stigmatized:

*(Clinician): “Some actually call me to bring their drugs to certain places. They don’t want to be coming to the facility always….. Their coming here regularly is raising questions in the community”*

Another perceived benefit reported by a minority of participants was that the early initiation of treatment under the new guideline has taking away the patients’ fear of dying:

*(Administrator):“So, that fear has been taking away. Immediately, I am tested, I am started on drugs”*

#### The perceived benefits of the Treat All guidelines to the health worker and the health system

The majority of the respondents reported that there have been significant benefits to the health workers and the health system since the introduction of the Treat All guidelines. For example, record keeping has improved because health workers were able to obtain complete records from patients because all activities relating to patients’ test and treatment were concluded on the same day:

*(Clinician): Okay, the promptness of it. And then the reduction in the stress on the part of the patient. And then even on the side of the health workers, there are advantages because it helps us get all the recordings the same day.*

*(Counsellor): “On our own side (health worker) it has increased our hope of recovery for the patient”*

A major perceived benefit to the health system was a reduction in perceived pre-treatment attrition. All respondents agreed that there has been a reduction in the initial patient loss between when patients tested HIV positive and when they started ART because there is essentially no more delays in starting treatment. However, there was a division of opinion on the attrition after patients started ART. The findings appeared to vary by treatment centre but no difference in opinion by health worker type. A majority of health workers interviewed in primary and secondary health centres perceived that there has been no change in lost to follow up among patients who have already started ART:

*(Clinician): “I wouldn’t say that has changed because most of the causes of lost to follow up in our environment is financial problems basically. So, I think it’s about the same”*

Whereas, respondents from tertiary health centres were of the opinion that lost to follow up has increased since the implementation of Treat All:

*(Clinician): “But, talk about lost to follow up, I think we have more lost to follow-ups since we started test and treat [Treat All]”*

Yet again, a minority of health workers in primary health setting perceived that there might actually be a reduction in lost to follow up among patients on ART since the commencement of Treat All:

*(Clinician):”There is no lost to follow up again because they already know they are on drugs (…) Lost to follow up is reduced to the barest minimum if there is any”*

### Perceived disadvantages of the Treat All guidelines

Respondents identified some drawbacks due to the implementation of the new Treat All guidelines at the patient and level of the health worker/health system. The following section described health workers perceived disadvantages of the treat all guidelines.

#### The perceived disadvantage of the Treat All guidelines to the patients

Health workers reported that a major success from the previous HIV treatment guidelines was a three weeks’ treatment preparation in which patients received an intensive education and counselling session, which stood out from other pre-treatment requirements as being useful and well-liked. The treatment preparatory class not only educated the patients but also created a platform for interaction between patients and health workers and patients with other patients. However, with the implementation of the new guidelines the treatment preparatory class is no longer available:

*(Administrator): “unlike the previous guidelines whereby you get to know your status and you are taken on a three weeks treatment preparatory class for you to get an understanding of all it takes to be positives and living with the virus. But, now your get tested immediately and you are started on drugs”(..) they might not really appreciate what it takes to be living with the virus”*

#### The perceived disadvantage of the Treat All guidelines to the health worker and the health system

The majority of respondents expressed concerns that the new guidelines have increased workload because under new guidelines standards, all PLHIV not currently receiving treatment must be offered treatment, and all newly-diagnosed PLHIV must start treatment immediately. The health workers explained that since the implementation of the Treat All guidelines, the number of patients eligible to start treatment has increased without a corresponding increase in the number of health workers providing health care. Consequently, the clinics have been crowded, increasing patient waiting time. Moreover, patients are often wary of coming to clinics if they perceive that they will spend a long time waiting.

*(Clinician): “Because the test and treat [Treat All guidelines] have increased the workload. So making the clinic a busy one for the clinicians”*

*(Clinician): “They are not waiting for you to waste their time anymore”*

### Reported patients’ response to Treat All guidelines

The majority of health workers elucidated that patients were generally happy with the change in the Treat All guidelines and were willing to start treatment early:

*(Administrator): “They are happy because many of them I come in contact with the first question they ask is where is the drug?”*

*(Counsellor): “There is a lot of enthusiasm now, patients now come and before you are done they are looking at you. They are expecting that they will be placed on treatment”*

Because Treat All was initially piloted in only 32 out of the 774 LGAs in Nigeria, respondents highlight instances where patients changed from health care providers who were yet to implement the new guidelines to those who had started implementing the guidelines. The primary reason for the change was because patients were not interested in wasting their time waiting before starting treatment:

*(Clinician): “we have seen patients changing from their places to come to us. Because, they heard that once you do the test today, you don’t have to waste time.”*

### Factors affecting guidelines use, perceived enablers

The factors identified by all of health workers interviewed as enablers were all health systems related factors. Health workers report that an important enabler in implementation of Treat All was training on the use of the guidelines. Participants explained that training gave the capacity to utilize the guidelines. Therefore, all health workers trained were willing to start using the guideline after receiving training:

*(Clinician): “They educated every person that participated actively in this program. You understand? And once they understood, it was easy for them to follow up”*

Participants further explained that there has been a variety of factors enabling the use of the Treat All guidelines at the various hospitals. The factors enabling guidelines use include the passion and interest of the health worker in caring for patients:

*(Clinician): “Okay, number one for me is the passion of the health workers*”

Another important enabler to the use of the guidelines was the availability of tools required to work, such as medications, test kits, job aids, and the written guidelines themselves in most clinics. Respondents further explained that health workers were likely to implement the Treat All guidelines in hospitals where the staff have sufficient supply of test kits and drugs. Furthermore, the availability of job aids and guidelines makes it easy for referencing and utilization of policy direction in the guidelines.

*(Administrator): “Hmmm, (…) the availability of the tools (…) they also brought job aids, It has helped (…) at least to refresh on the knowledge from the training”*

*(Clinician): “Because if we didn’t have drugs that will mean the clients will come and we will not be able to start their treatment”*

Respondents further explained that good management support and regular government supervisory visits were added motivation for implementing the guidelines.

*(Administrator): “since the supervisory visit is regular, they give you an update or give you a step-down and say this is how it should be or do it this way”*

### Factors affecting guidelines use, perceived barriers

The factors perceived as barriers were outlined into two broad categories, at the levels of the patient and the health system. The following sections describe the barriers by patient and health system levels.

#### Factors serving as barriers at the patient level

Participants highlighted several factors that made it difficult to implement the Treat All guidelines. However, the most important barrier that cut across nearly all study sites was poverty. The patient’s inability to pay for transportation to site or for baseline investigations was a major barrier to the implementation of the test and treat guidelines.

*(Clinician): “A patient comes in and obviously you just see anemia, why are you not eating what was recommended? They will say they don’t have money”*

*(Counsellor): “so out of about five of them I saw this morning only one assured me that he will come to the meeting the other one said mummy no transport”*

#### Factors serving as barriers on the health workers/health system level

A minority of respondents identified stock out of test kits, which occurs occasionally, as another barrier. The stock out of test kits makes it impossible to test new patients for HIV in affected hospitals. Some health workers also cited a lack of sufficient copies of guidelines for staff review as a barrier to effective implementation.

*(Clinician): Though sometimes we run short of test kits. It is important that everything we need to work is available. Sometimes there we [are] short of supplies.*

*(Clinician): “Yes, another barrier is if you don’t have enough copies of the guideline for everybody (health workers) to have. In this facility doctors change frequently so the two or three copies they brought is not enough”*

Additionally, a lack of a dedicated space to provide treatment for tuberculosis, which is considered the most common opportunistic infection among PLHIV in Nigeria, and more generally a lack of integration of TB care with HIV care, may constitute a barrier to ensuring comprehensive care among PLHIV in some clinics.

*(Administrator): We have a challenge with the DOT clinic, our TB/HIV integration is not doing so well.*

Furthermore, the level of training may be a barrier because younger doctors do not have enough information about the guidelines and may require more training to come up to speed about the change.

*(Clinician): “The level of training may be a barrier, we have younger doctors, house officers, youth corpers, they are here for just one year”*

Finally, respondents explained that inadequate human resources, specifically, low numbers of health staff, is a major barrier to implementation of the new guidelines. As the number of patients continue to increase, respondents felt that there needs to be a corresponding increase in number of health workers to ensure quality of care is not compromised.

*(Clinician): “Then another barrier is manpower. Because the test and treat [Treat All] has increased the workload. So making the clinic a busy one for the clinicians”*

### Other findings

The following section outlines incidental findings that were outside of the primary purpose of our study, but were nonetheless important and of value.

#### Factors contributing to lost to follow among patients on ART

Although not the main focus of the project, through the discussions participants described what they perceived to be factors associated with loss to follow-up. Health workers reported that there were multiple issues affecting lost to follow up among patients initiated on ART. Factors contributing to lost to follow up include financial challenges, making the patients who live far away from clinics unable to afford transportation to clinics or pay for investigations not covered by the program:

*(Clinician): “if we hammer on the need for an investigation that the patient needs and cannot afford, the patient might become afraid and not show up anymore (……) Another thing is proximity; some may be eager to start but won’t tell you that they stay far away from the facility (…..) we say the patient is lost to follow up not knowing that the patient actually self-transferred”*

Other factors reported by health workers contributing to attrition include religious/traditional belief, stigma and denial, lack of education, and self-transfer:

*(Administrator): “Experience of lost to follow up is due to several factors. Number one is transportation, number two is location. Some of them choose to go on their own, self-transfer”*

*(Clinician): “Religious rulers are affecting (…) because we almost lost one patient last year. Because according to her the pastor prayed for her and told her that all is well that she should stop the drugs”*

*(Clinician): “Because most educated people agree, they will take it and you will see them coming back”*

Respondents report that patients were also not satisfied with the repeated hospital visits without receiving medications all because they were not eligible to start treatment by the standards of the previous guidelines. Consequently, some indigent patients stop coming to the hospital and are lost to follow up after many repeated unfruitful visits to the clinic:

*(Clinician): “But you know sometimes when you come today, come tomorrow, (…) the patients, some don’t have the means of coming back”*

#### Late responders

A minority of respondents reported that although most patients were perceived to be willing to start treatment early, there were some patients who were not willing to start treatment early. Health workers perceive that this category of patients is composed of ‘late responders’ who need more time to make up their minds to start ART. When informed that they are HIV positive they have a strong negative reaction. Participants further explained that during counseling this kind of patients can be identified by the following characteristics: denial, self-stigmatization, depression, and discouragement. These patients are not enthusiastic about starting treatment and some may refuse to give contact details and therefore cannot be contacted if they default in attending their clinical appointment:

*(Administrator): “those that want to live in denial and want to say(…) ha! I don’t have the virus! Because they look healthy, they feel that (…) I am not sick, I am doing fine so why should I go on therapy”*

If started on treatment immediately, these “late responders” fail to return for further follow-up visits: *(Clinician): “And by the time I finish everything, she was given the drug for two weeks but because there was no phone number no place to trace her. Because I just couldn’t trace her, I tried…”*

The “late responders” may require several follow up counseling sessions before they eventually make up their minds to be committed to treatment:

*(Counsellor): “But, some of them (…) ha!(…) I don’t like drugs. You can’t just push me into drugs immediately… give me some time”*

### How to improve perceived barriers

Interviewed participants provided some very important ideas for improving on some of the barriers identified during the study, in order to improve and scale up implementation of the Treat All guidelines nationwide. A major recommendation was for the government to introduce some form of financial assistance scheme to support patients who could not afford to transport themselves to clinics or pay for required laboratory investigations.

*(Counsellor): “Some of them have problem of coming here with their TP (….) [Transportation] … for some of them feeding problems and all that. (……). At each of their visit if we can give a little thing, a little token to help them so they can come back next time for their visit”*

The Treat All guidelines pilot implementation team could help improve and scale up implementation through advocacy to the government on the need to provide additional clinical staff to support the growing number of patients at the clinics. The implementation team can also advocate for laboratory staff at the laboratory to reduce processing time for the additional viral load tests.

*(Clinician): “I think there is still need for more advocacy for some clinicians”*

*(Administrator) “it is to involve the lab. We need to get them together and tell them the implications of not starting the patients on treatment”*

Another area that a government intervention will help with is in improved support to assist TB/HIV integration to ensure comprehensive care for PLHIV.

*(Administrator): So, I want to make a suggestion, if we can find a way to help to revitalize the DOT section of the hospital*

Another area is ensuring more government support to sites through supervision to the hospitals and provision of sufficient copies of the guidelines to all HIV clinics including more concise pocket versions of the guidelines.

*(Clinician): “Yes, another barrier is if you don’t have enough copies of the guideline for everybody. There could be pocket versions of the guidelines that will make it easy to carry about”*

Lastly, hospitals could implement a more differentiated service delivery approach that takes services into the communities to decongest the clinics.

*(Clinician): it has helped to decongest the clinic. For example, some of the patients don’t need to come to the clinic again they just go to the community pharmacy to pick up their drugs*

## Discussion

The implementation of Treat All guidelines in Nigeria was perceived to have significantly changed clinical practice for care of PLHIV, bringing about reduction in delays before treatment start, reduction in stress and increase in hope among patients. Additionally, because all events leading to treatment initiation are completed on the same day, there are more complete records for patient follow-up. Empirical studies have been used to evaluate outcomes following changes in treatment guidelines. A quantitative study conducted in South Africa, evaluating the impact of a Treat All approach, used a mathematical modelling technique and partial differential equations [20]. The study reported that there was sufficient evidence demonstrating that doing more testing for HIV, followed by immediate treatment of persons found HIV positive, could ‘eradicate’ the HIV epidemic and reduce the cost for HIV medication in the long run [20]. Additionally, a randomized control trial conducted at 215 sites in 35 countries among 2,359 HIV infected individuals assessed outcomes using the Treat All approach [21]. The study reported that implementing the Treat All guidelines led to better outcomes than delaying treatment [21]. Furthermore, a multicentre retrospective cohort study conducted in four hospitals in Kenya analysed outcomes of pregnancies amongst 1,365 HIV+ pregnant women who received ART intervention using the Treat All approach. The study reported a 50% reduction in HIV transmission from mothers to their children due to the Treat All approach [22]. The reports from these empirical studies supported the perception of health workers in this study of better patient outcomes using the Treat All guidelines.

Among health worker respondents, the perception was that the majority of patients were happy with the guideline change because of a perception of good health benefits when treatment is started early. Similarly, health workers were also willing to implement the new guidelines for related reasons. The study finding was consistent with reports by Nzinga *et al* and Odhiambo *et al*. Both studies observed that knowledge of the benefits from implementing guidelines was a major motivational factor to utilizing guidelines [15,23]. The study results suggest that new public health initiatives such as guidelines or policy changes will require significant health worker and patient education to increase the chances of successful implementation and improved patient outcomes. Specifically in the Nigerian context, health workers perceive that the expansion of Treat All nationally will require patient education about the health benefits of starting treatment early. In contrast, the category of patients referred to as “late responders” were unwilling to start treatment early because they were still in denial and therefore not yet convinced of any benefit of starting treatment early. Obligating these patients to start treatment before they are ready may lead to these patients being lost to follow up. This finding reaffirms the need for screening for and establishing patient readiness before starting ART [24]. Thus, though the expansion of Treat All nationally in Nigeria will entail early initiation of the vast majority of PLHIV on ART, there should be support for identifying and working with PLHIV who may need longer to be ready to initiate ART, in order to avoid attrition.

Health workers perceived that the introduction of the Treat All guidelines led to a reduction in mortality and improvements in viral suppression and patient overall health. This perception was consistent with findings previously reported in the literature [21,22,25]. However, there was a perception that the Treat All guidelines alone cannot resolve lost to follow up in patients on ART. Health workers perceived that there were multiple factors responsible for lost to follow up among patients that started ART immediately, topmost among them was financial difficulties, making it challenging to afford transport to clinic and payment for investigations. This finding was consistent with those previously reported [26-28]. Maskew *et al*, in their study in South Africa, conducted among 182 HIV positive patients who missed follow up appointments, and contacted through telephone calls, reported that a majority of their patients cited financial difficulties and inability to pay for transportation costs as major reasons for missed appointments [26]. Consequently, there are no guarantees that starting treatment early may lead to a reduction in lost to follow up. The implication of this finding is that program managers must adopt strategies that empower the clients to keep their clinic appointments and treatment approaches that take services closer to communities to reduce lost to follow up [29-31].

Factors previously identified as enablers of guidelines use included collaboration between facilities, peer support, and provider characteristics such as higher level of education, commitment, and knowledge of guidelines [32]. The major factors identified in the literature that serve as barriers to treatment guidelines implementation included human resource gaps, lengthy guidelines, inadequate supplies of medicines, and commodities [32]. This study found that training, availability of tools, government supervisory visits, hospital management support and health workers passion and interest enabled guidelines use. Identified barriers in this study included patient poverty, inadequate human resources, and stock-outs of tools, guidelines, and test kits. Thus, the identified enablers and barriers to guidelines use in this study were comparable to those previously reported [32]. The implication of these findings is that program managers must ensure that clinics have staff strength adequate for patient load, ensure adequate supervisory visits, provide adequate supplies of test kits and guidelines and implement innovative approaches that decongest clinics in order to improve the impact of the implementation of the Treat All guidelines. Each of these will need to be addressed as Nigeria scales up Treat All nationally.

Focusing specifically on patient poverty as a main barrier to Treat All implementation, the study revealed that patients’ lack of capacity to pay for baseline investigation or transport services to clinics limited their ability to access treatment services. The results re-affirms the concept of a wider set of determinants for health and the important effect of socioeconomic factors in determining the health of individuals [33-35].

The study demonstrates that the presence of social support systems can influence an individual’s health-seeking behaviours [36]. The study results suggest that to improve retention among the higher number of PLHIV on ART inherent in the Treat All approach, HIV programs in Nigeria may need to incorporate social support safety nets to help indigent patients.

Health worker respondents perceived that the challenges of patient access could be mitigated by implementation of patient empowerment schemes that increase patients’ skills and empower them economically to provide for themselves and their households. An economically empowering scheme implemented in Kenya enabled HIV-positive persons through capacity building and provision of soft loans to become financially sufficient [37]. A similar approach in Nigeria may contribute to improved attendance at clinics as Treat All expands nationally. From the perspective of the health workers, patients still come from distant communities to access HIV care and treatment in big cities. This finding suggests that implementing community-based HIV treatment models that take services into communities using various approaches such as mobile clinics, community adherence clubs and community drug dispensing outlets may reduce the need for patients to travel far and further improve adherence to treatment [29,31,38].

Focusing specifically on inadequate health staff as a main barrier to Treat All implementation, the government could resolve this challenge by scaling up a variety of approaches including the use of six-month prescriptions that give stable patients up to six months’ drug refills at a time, drastically reducing the frequency for clinic visits [39]. Moreover, the current treatment guidelines in Nigeria supports the use of multi-month scripting [8]. Additionally, Nigeria already has a policy on task shifting in HIV care and treatment, and reports suggests that these approaches are currently being implemented and are yielding good results in maintaining PLHIV on treatment [40-42]. Therefore, scaling up approaches such as task shifting and task sharing that build the capacity of nurses to provide clinical care and allow nurse prescription of ART could further expand the existing human resource capacity for HIV services in Nigeria [41,42].

The strengths of the study included the rigorous process to ensure conformity with established qualitative research and ethical standards. Additionally, the use of the interview guide and appropriate probes ensured that all participant were measured using similar standards [43], thus reducing measurement and social desirability biases. To mitigate the role of positionality, the researchers ensured that their views were bracketed by identifying their own preconceived assumptions and setting them aside during the interview process [44,45]. A limitation of the study was the authors’ inability to triangulate health workers’ perceptions to patients’ reactions of the guideline changes because patients’ perspectives were not collected. A qualitative design that obtains the perspectives of the patients themselves could have addressed this challenge and is a potential future area of research [46,47]. Additionally, health worker perceptions of patient outcomes were not matched to actual patient outcomes, limiting the ability of this study to comment on correlation of guidelines changes and any change in patient outcomes of interest, including viral suppression and mortality [48]. Another major limitation of the study was the limited cultural context of its findings since the study was conducted in one state in the country limiting the transferability of its findings [49].

## Conclusion

According to participating health workers, the introduction of the Treat All guidelines in Nigeria improved access to skilled HIV care and treatment services for PLHIV and were perceived to improve patients’ overall health. Specific areas perceived to have improved after guidelines change included viral suppression, mortality, and waiting time at clinics. Factors enabling the implementation of the treat all guidelines were training of health workers on guideline use, the passion and interest of health workers to see patients get better, and the availability of tools, government supervisory visits and good hospital management support. The identified barriers to successful guideline implementation were poverty, inadequate human resource at clinics, and non-availability of tools, guidelines and test kits at some hospitals. Study findings suggest that the effective implementation and national scale up of the Treat All guidelines, with the intention of improving PLHIV clinical outcomes, could be further enhanced if the government provided support for indigent patients, more health worker staff, adequate supervisory visits, adequate supplies of test kits, and guidelines. In addition, implementing innovative approaches that decongest clinics by providing more services in communities would enhance effective implementation of the Treat All guidelines. In these ways, the Treat All approach can be effectively expanded nationally, thus giving all PLHIV in Nigeria timely access to comprehensive HIV care and treatment.

## Acknowledgements

We will like to appreciate the staff of the Federal Ministry of Health, institute of human virology and the sites in which the study was conducted for their support during interviews. Our most heartfelt gratitude also goes to Mr. Folorunsho Eyitayo for his commitment and support during the study. Lastly, we would like to express our most profound gratitude to Dr. Constance Shumba for her dedicated support to this work.

## Disclaimer

The findings and conclusions in this manuscript are those of the authors and do not necessarily represent the views of the Funding agencies. Use of trade names is for identification only and does not imply endorsement by the U.S. Centers for Disease Control and Prevention or the U.S. Department of Health and Human Services.

## References

1. WHO (2017) HIV Treatment and Care. Treat All: Policy adoption and implementation status in countries (Factsheet). Geneva, Switzerland: WHO. pp. 4.

2. UNAIDS (2016) Global AIDS update. Geneva: UNAIDS.

3. WHO (2017) HIV/AIDS Factsheet. WHO Publications. Geneva: Switzerland.

4. UNGASS (2016) Political Declaration on HIV and AIDS: On the Fast Track to Accelerating the Fight against HIV and to Ending the AIDS Epidemic by 2030. In: UNAIDS, editor. Resolution adopted by the General Assembly on 8 June 2016. Geneva: UNAIDS.

5. bank W (2018) Country population (Nigeria). All Countries and Economies (Population, Total). Washington DC: World bank

6. UNAIDS (2018) Country Factsheets: Nigeria. HIV and AIDS Estimates. Geneva: UNAIDS.

7. PEPFAR Nigeria (2016) Strategic Direction for 2016. Nigeria Country Operational Plan. Abuja, Nigeria: United States Embassy.

8. FMOH (2016) National guidelines for HIV Prevention Treatment and Care. In: National AIDS & STI Control Programme, editor. Abuja: FMOH.

9. Adeloye D, David RA, Olaogun AA, Auta A, Adesokan A, et al. (2017) Health workforce and governance: the crisis in Nigeria. Hum Resour Health 15: 32.

10. Boyce C, Neale P (2006) Conducting in-depth interviews: A Guide for Designing and Conducting In-Depth Interviews for Evaluation Input. Monitoring and Evaluation Watertown, Massachusetts: Pathfinder international.

11. Statistics NBo (2018) Population of Abuja. Population of Nigeria. Abuja: National Bureau of Statistics.

12. Federal Ministry of Health (2015) Sentinel Survey among Pregnant Women Attending Antenatal Clinics in Nigeria. In: National AIDS/STI Control Programme, editor. Abuja: Federal Ministry of Health.

13. Federal Ministry of Health (2017) Fiscal Year 2017 Quarter three data. Washington DC: PEPFAR

14. Ayres L (2007) Qualitative research proposals--part III: sampling and data collection. J Wound Ostomy Continence Nurs 34: 242–244.

15. Nzinga J, Mbindyo P, Mbaabu L, Warira A, English M (2009) Documenting the experiences of health workers expected to implement guidelines during an intervention study in Kenyan hospitals. Implement Sci 4: 44.

16. Braun V, Clarke V (2006) Using thematic analysis in psychology. Qualitative Research in Psychology 3: 77–101.

17. Miller R (2013) Inductive Coding. Development of a Proposal for a Technology Assessment Report. Edinburgh: University of Edinburgh,.

18. Shenton AK (2004) Strategies for ensuring trustworthiness in qualitative research projects. Education for Information 22: 63–75.

19. Allmark PJ, Boote J, Chambers E, Clarke A, Mcdonnell A, et al. (2009) Ethical issues in the use of in-depth interviews: literature review and discussion. Research ethics review 5: 48–54.

20. Dodd PJ, Garnett GP, Hallett TB (2010) Examining the promise of HIV elimination by ‘test and treat’ in hyperendemic settings. AIDS 24: 729–735.

21. Group ISS, Lundgren JD, Babiker AG, Gordin F, Emery S, et al. (2015) Initiation of Antiretroviral Therapy in Early Asymptomatic HIV Infection. N Engl J Med 373: 795–807.

22. Finocchario-Kessler S, Clark KF, Khamadi S, Gautney BJ, Okoth V, et al. (2016) Progress Toward Eliminating Mother to Child Transmission of HIV in Kenya: Review of Treatment Guideline Uptake and Pediatric Transmission at Four Government Hospitals Between 2010 and 2012. AIDS Behav 20: 2602–2611.

23. Odhiambo GO, Musuva RM, Odiere MR, Mwinzi PN (2016) Experiences and perspectives of community health workers from implementing treatment for schistosomiasis using the community directed intervention strategy in an informal settlement in Kisumu City, western Kenya. BMC Public Health 16: 986.

24. Rosen S, Fox MP, Larson BA, Brennan AT, Maskew M, et al. (2017) Simplified clinical algorithm for identifying patients eligible for immediate initiation of antiretroviral therapy for HIV (SLATE): protocol for a randomised evaluation. BMJ Open 7: e016340.

25. Dodd PJ, Garnett GP, Hallett TB (2010) Examining the promise of HIV elimination by ‘test and treat’ in hyperendemic settings. AIDS 24: 729–735.

26. Maskew M, MacPhail P, Menezes C, Rubel D (2007) Lost to follow up: contributing factors and challenges in South African patients on antiretroviral therapy. S Afr Med J 97: 853–857.

27. Megerso A, Garoma S, Eticha T, Workineh T, Daba S, et al. (2016) Predictors of loss to follow-up in antiretroviral treatment for adult patients in the Oromia region, Ethiopia. HIV AIDS (Auckl) 8: 83–92.

28. Cattamanchi A, Miller CR, Tapley A, Haguma P, Ochom E, et al. (2015) Health worker perspectives on barriers to delivery of routine tuberculosis diagnostic evaluation services in Uganda: a qualitative study to guide clinic-based interventions. BMC Health Serv Res 15: 10.

29. Decroo T, Telfer B, Dores CD, White RA, Santos ND, et al. (2017) Effect of Community ART Groups on retention-in-care among patients on ART in Tete Province, Mozambique: a cohort study. BMJ Open 7: e016800.

30. Okoboi S, Ding E, Persuad S, Wangisi J, Birungi J, et al. (2015) Community-based ART distribution system can effectively facilitate long-term program retention and low-rates of death and virologic failure in rural Uganda. AIDS Res Ther 12: 37.

31. Decroo T, Koole O, Remartinez D, dos Santos N, Dezembro S, et al. (2014) Four-year retention and risk factors for attrition among members of community ART groups in Tete, Mozambique. Trop Med Int Health 19: 514–521.

32. Mwangome MN, Geubbels E, Wringe A, Todd J, Klatser P, et al. (2017) A qualitative study of the determinants of HIV guidelines implementation in two south-eastern districts of Tanzania. Health Policy Plan 32: 825–834.

33. Whitehead M (2007) A typology of actions to tackle social inequalities in health. J Epidemiol Community Health 61: 473–478.

34. Dyson YD, Mobley Y, Harris G, Randolph SD (2018) Using the Social-Ecological Model of HIV Prevention to Explore HIV Testing Behaviors of Young Black College Women. J Assoc Nurses AIDS Care 29: 53–59.

35. Drew S, Lavy C, Gooberman-Hill R (2016) What factors affect patient access and engagement with clubfoot treatment in low-and middle-income countries? Meta-synthesis of existing qualitative studies using a social ecological model. Trop Med Int Health 21: 570–589.

36. Frieden TR (2010) A framework for public health action: the health impact pyramid. Am J Public Health 100: 590–595.

37. Weiser SD, Bukusi EA, Steinfeld RL, Frongillo EA, Weke E, et al. (2015) Shamba Maisha: randomized controlled trial of an agricultural and finance intervention to improve HIV health outcomes. AIDS 29: 1889–1894.

38. Jobarteh K, Shiraishi RW, Malimane I, Samo Gudo P, Decroo T, et al. (2016) Community ART Support Groups in Mozambique: The Potential of Patients as Partners in Care. PLoS One 11: e0166444.

39. Fatti G, Ngorima-Mabhena N, Chirowa F, Chirwa B, Takarinda K, et al. (2018) The effectiveness and cost-effectiveness of 3- vs. 6-monthly dispensing of antiretroviral treatment (ART) for stable HIV patients in community ART-refill groups in Zimbabwe: study protocol for a pragmatic, cluster-randomized trial. Trials 19: 79.

40. Umar NA, Hajara MJ, Khalifa M (2011) Reduction of client waiting time using task shifting in an anti-retroviral clinic at Specialist Hospital Bauchi, Nigeria. J Public Health Afr 2: e2.

41. Rustagi AS, Manjate RM, Gloyd S, John-Stewart G, Micek M, et al. (2015) Perspectives of key stakeholders regarding task shifting of care for HIV patients in Mozambique: a qualitative interview-based study with Ministry of Health leaders, clinicians, and donors. Hum Resour Health 13: 18.

42. Iwu EN, Holzemer WL (2014) Task shifting of HIV management from doctors to nurses in Africa: clinical outcomes and evidence on nurse self-efficacy and job satisfaction. AIDS Care 26: 42–52.

43. Kvale S (1994) Ten standard objections to qualitative research interviews. Journal of Phenomenological Psychology 25: 147–173.

44. Sorsa MA, Kiikkala I, Astedt-Kurki P (2015) Bracketing as a skill in conducting unstructured qualitative interviews. Nurse Res 22: 8–12.

45. Fischer CT (2009) Bracketing in qualitative research: conceptual and practical matters. Psychother Res 19: 583–590.

46. Muleme J, Kankya C, Ssempebwa JC, Mazeri S, Muwonge A (2017) A Framework for Integrating Qualitative and Quantitative Data in Knowledge, Attitude, and Practice Studies: A Case Study of Pesticide Usage in Eastern Uganda. Front Public Health 5: 318.

47. Wong LP (2008) Focus group discussion: a tool for health and medical research. Singapore Med J 49: 256-260; quiz 261.

48. Dalhatu I, Onotu D, Odafe S, Abiri O, Debem H, et al. (2016) Outcomes of Nigeria’s HIV/AIDS Treatment Program for Patients Initiated on Antiretroviral Treatment between 2004-2012. PLoS One 11: e0165528.

49. Green J, Thorogood N (2014) Developing Qualitative Analysis. Qualitative Methods for Health Research. Thosand Oaks, CA: Sage Publications Inc. pp. 250–253.

